# Seasonal changes in the physiology and metabolism of grapevine perennating buds

**DOI:** 10.1101/2025.06.17.660214

**Authors:** Santiago Signorelli, Dina Hermawaty, Regina Feil, Camila Couture, John A. Considine, John E. Lunn, Michael J. Considine

**Author notes:** Correspondence: Santiago Signorelli.

## Abstract

Grapevine (*Vitis vinifera L.*) buds undergo seasonal dormancy to survive unfavourable conditions and synchronize growth with environmental cues. While dormancy transitions have been widely studied in temperate woody perennials, the physiological and metabolic dynamics underlying these transitions in grapevine remain poorly understood. Our study investigates seasonal changes in the physiology and metabolism of perennating buds in *V. vinifera* cv. Cabernet Sauvignon, focusing on dormancy depth, respiration, sugar metabolism, and cell cycle activity. We identified three distinct phases of bud quiescence: (i) para-dormancy (early summer), characterized by active metabolism and high levels of tricarboxylic acid cycle intermediates, shikimate, and myo-inositol; (ii) endo-dormancy (late summer to autumn), where dormancy depth peaked in late summer and was marked by reduced respiration and bud water content, and a sharp decline in hexose levels with a concomitant increase in raffinose; and (iii) eco-dormancy (winter to spring), featuring increased respiration, sugar mobilization (notably sucrose, glucose, and trehalose), and reactivation of cell division, with a shift of cells into the G2 phase. Just prior to bud burst, we observed a significant accumulation of sugar-phosphates, providing evidence that supports their role in promoting bud burst also in grapevine. Our findings provide new insights into the biochemical and physiological regulation of bud dormancy and bud burst, contributing to a deeper understanding of dormancy transitions in perennial crops.

## Introduction

Grapevine (*Vitis vinifera* L.) is the most economically important fruit crop worldwide, grown and commercially traded between over 190 countries (faostat.org). Grapevine was domesticated as a deciduous perennial (Terral *et al*., 2010), however it displays wide plasticity to climate when grown in warmer, lower latitude climates (Camargo *et al*., 2012; OIV, 2017; Velappan *et al*., 2022*c*). Combined with increasing land pressure for arable land, and long-term climate change, understanding the eco-physiology of grapevine through the whole year is fundamentally important.

The vitality of the perennating bud (syn. latent bud, N+2; Lavee and May, 1997) is vital to sustainable production, as it bears the vegetative and reproductive initials that form the next cycle of production following growth arrest and bud burst (Lavee and May, 1997). The bud develops under quiescence over a period of nearly 12 months or four seasons, and yet our knowledge of the eco-physiology, and metabolic control of the bud is largely restricted to winter and spring (De Rosa *et al*., 2021; Velappan *et al*., 2022a). Hence, a considerable need exists for a more integrated understanding of the development and vitality of the bud throughout its lifecycle. This includes the transitions into and out of dormancy in the strict sense.

The major seasonal cues of light and temperature affect the activity of axillary buds by influencing energy and hormone functions. In non-dormant grapevine buds subjected to forcing conditions, light was shown to induce a photomorphogenic response and accelerated bud burst rates (Signorelli *et al*., 2018, 2020a). In other woody perennial species such as *Rosa* spp., light is an obligate requirement for bud growth, where it was shown to activate sugar mobilization and induce the expression of vascular invertase enzymes (Girault *et al*., 2010; Rabot *et al*., 2012). Likewise, the sorghum (*Sorghum bicolor* L.) mutant for phytochrome B (phyB) was shown to be unable to fully develop axillary buds, and this was correlated with lower expression of SUGARS WILL EVENTUALLY BE TRANSPORTED (SWEET) sugar transporters and cell wall invertase, suggesting that light perception is key to promote a correct apoplastic supply of sugars that promotes bud development (Kebrom and Mullet, 2016). Moreover, sucrose was shown to augment the effect of light on bud outgrowth (Rabot *et al*., 2012). This effect was also observed when applying non-metabolizable sucrose analogues but not when applying mannitol (Rabot *et al*., 2012; Barbier *et al*., 2015). These observations suggest that, although sucrose could have a metabolic effect, its impact was primarily via sugar signalling.

In the last decade, several fundamental studies have shown the importance of sugars in metabolic and hormone control of quiescence and bud outgrowth in annual herbaceous plants (Considine and Foyer, 2023); however, our knowledge about these processes in woody perennials is much more limited. Defoliation, which is known to inhibit bud outgrowth, was shown to induce sucrose-starvation gene markers in axillary buds of sorghum within 24 hours (Kebrom and Mullet, 2015), whereas shoot tip removal rapidly induced sugars, resulting in bud outgrowth of pea (*Pisum sativum* L.) (Mason *et al*., 2014). Moreover, decapitation of pea shoots resulted in a rapid increase of trehalose 6-phosphate (Tre6P), a sucrose signalling metabolite, and amino acid levels, while decreasing phosphoenolpyruvate and 2-oxoglutarate levels, resulting in bud outgrowth (Fichtner *et al*., 2017). Further, the heterologous expression of Tre6P phosphatase in Arabidopsis (*A. thaliana* (L.) Heynh.) resulted in lower Tre6P in axillary buds and delayed bud outgrowth and decreased branching (Fichtner et al., 2021). Recently, the sugar-responsive S1-basic-leucine-zipper (S1-bZIP) transcription factor was shown to be involved in these processes by regulating the carbohydrate partitioning from leaves to the apical meristem, via the control of the expression of SWEET sugar transporters (Kreisz *et al*., 2024). The absence of S1-bZIP function results in a greater lateral growth development (Kreisz *et al*., 2024). Moreover, mechanistic relationships have been demonstrated between light signalling, glucose and the TARGET OF RAPAMYCIN (TOR) kinase to explain meristematic activity of Arabidopsis (Xiong *et al*., 2013a; Pfeiffer *et al*., 2016; Pereyra *et al*., 2023). Thus, light and sugars are integrated signals controlling meristematic activity and bud outgrowth.

Despite the advanced understanding of quiescence/ activity cycles in arabidopsis and other herbaceous annuals, the seasonal cycles of quiescence and dormancy in grapevine represent a considerably more complex ecological function. Present knowledge of the dynamics and functions of sugars and other organic metabolites in grapevine seasonality is sparse. Here, we address this gap, by examining how the dynamics of sugars and organic acids relate to physiological, metabolic and climate variables during the bud quiescence and dormancy cycles of grapevine from mid-summer through early spring. These data provide a benchmark to advance our understanding of the role of metabolic control in the perception and transduction of seasonality in important perennial species.

## Methods

Unless otherwise stated all chemicals were analytical grade from Sigma Aldrich (NSW, Australia; www.sigmaaldrich.com).

### Plant material and growth conditions

Canes of *Vitis vinifera* (L.) cv. Cabernet Sauvignon containing buds from node 3 to 12 were collected from a vineyard in Margaret River (33°47′S 115°02′E), WA, Australia. The sampling dates were 5 January, 15 February, 4 April, 10 May, 28 June, 9 August and 20 September. At the earliest sampling time (January), all basal buds up to node >15 were mature and lignified. These canes were transported to the laboratory and immediately analysed or planted for time-course experiments.

### Depth of dormancy

For each sampling date, two replicates of 50 single-node cuttings were planted in pots containing a soil mix (fine composted pine bark, coco peat, and brown river sand in the ratio of 2.5:1:1.5). As a positive control for viability, the same number of single-node cuttings were immersed in 1.25% (v/v) hydrogen cyanamide (HC) (0.31 M) in water for 30 seconds before planting. Explants were grown in a controlled environment chamber under dark-light (12:12 h) conditions at 23 °C and the irradiance, provided by light-emitting diodes, was between 200-400 µmol photons m^-2^ s^-1^. Bud burst was recorded at least three times a week, and the depth of dormancy was estimated as the time required to reach 50 % of bud burst (BB50). Bud burst was scored when a green leaf tip became visible, i.e. at stage EL4 according to the modified Eichorn-Lorenz scale (Coombe, 2004). At the end of the experiment, all remaining buds were dissected and assessed for necrosis, and corrections were made to the cumulative bud break based on the number of healthy buds.

### Weather data

The weather data were obtained from Moss Wood vineyard in Margaret River. The average daily mean air temperatures were calculated from hourly air temperatures using the R statistical package (R Development Core Team, 2014) and plotted using ggplot2 package of R (Wickham, 2009) as scatter dot plots fitted with a quadratic spline with degree of freedom (df)=4 and degree=2. For chilling model we used the Daily Positive Utah Chill Unit (DPCU) model, which is a modified version of the Utah model suggested to accurately predict chill accumulation in mild winter regions (Linsley-Noakes *et al*., 1995). This model records chill as Positive Chill Units (PCU) and considers optimal chilling to occur at temperatures between 2.5°C and 9.2°C, and nil chilling at temperatures less than 1.5°C or greater than 12.5. Hence, 1h at 2.4°C < T ≤ 9.1°C equals 1PCU, 1h at 1.4°C < T ≤ 2.4°C or 9.1°C < T ≤ 12.4°C equals 0.5PCU and 1h at T ≤ 1.4°C or T ≥ 12.4°C equals 0PCU. The cumulative chilling hour or chill unit was estimated for each day from the hourly mean air temperature calculated and represented as a line plot using R package (R Development Core Team, 2014) and ggplot2 package of R (Wickham, 2009) respectively.

### CO_2_ release and O_2_ consumption

To determine the CO_2_ release, four pools of eight buds each were placed in an insect respiration chamber (6400-89; Li-COR, Lincoln, NB, USA; www.licor.com) attached to a Li-6400XT portable gas exchange system. The measurements were performed in complete darkness, at 23 °C, in CO_2_-controlled air (380 µmol CO_2_ mol^-1^ air) with 100 µmol s^-1^ air flow, at 55–75 % relative humidity.

Oxygen consumption was determined in four pools of eight buds each. The buds were placed in a 4 mL micro-respiration chamber and the O_2_ content was monitored for at least 10 min using an O_2_ microsensor (Clark-type, OX-25; Unisense, Aarhus, Denmark). Measurements were performed at 23 °C in complete darkness and the micro-respiration chamber was kept under water to prevent sudden changes in temperature. The volume of the buds was determined using a 25 mL density bottle, where water was used, and the volume was measured by determining the mg of water was displaced by the bud. This information was necessary to determine for each replicate the final volume of air inside the chamber.

For both CO_2_ release and O_2_ consumption, the buds were dried in an oven for 72 h at 70 °C to determine the dry weight (DW) and the respiration rates were referred to the DW of the buds. Respiratory quotient (RQ) values cannot be deduced directly from these data because the conditions of measurements were different for CO_2_ and O_2_. Therefore, we calculated a relative RQ (RRQ) in which the RQ was calculated for each sampling date, and it was divided by the average of the RQ, to normalize the average to 1. Hence the higher values indicate a more anaerobic metabolism, while the lower values indicate a more aerobic metabolism, assuming that carbohydrates were the predominant respiratory substrate.

### Bud water content

Three sets of ten buds each were used to determine the moisture content. The buds were dissected from the cane and the fresh weight was recorded. Then, they were desiccated during 96 h in an oven at 70 °C and the dry weight was determined. The moisture content was calculated as follows:

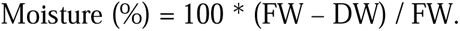

### Internal pO_2_ profiles

The internal oxygen was measured using an oxygen microsensor (Clark-type) with a 25 µm tip (OX-25; Unisense A/S, Aarhus, Denmark). The calibration of the sensor and measurements were done using the Sensortrace Suite software (https://unisense.com). For calibration at 0 kPa oxygen, the sensor was flushed with N_2_ until stabilization, and for air-saturated O_2_ concentration, an aquarium pump was used to flush the sensor with atmospheric air (21% O_2_). The buds were removed from the cane and stood on a flat surface and then the electrode was electronically inserted from the top to reach the centre of the meristem (Shaw *et al*., 2017). The depth of the path was about 2200 µm, depending on the size of the bud, with steps of 35 µm and three measurements per step were performed. At least four buds were used for each sampling date.

### Apoplastic connection

Eosin Y and Acid fuchsin solutions were prepared by dissolving 0.5 g of the respective dye in 0.01 M potassium phosphate buffer pH 6.0. The solutions were filtered through GF/A filter paper and then through a 0.2-µm membrane filter. Eight single-node cuttings for each sampling date were cut 1.5 cm below the base of the bud and immediately transferred to 25-mL plastic tubes containing 2 mL of Eosin Y or Acid fuchsin solutions (four replicates each), in a way that the cane was immersed but the bud was not submerged. The incubation time in the solution was 4 h. A razor blade was used to cut the cane in the longitudinal plane and the images were taken using a camera coupled to a magnifying glass.

### Cell cycle status

Intact nuclei were isolated from fresh buds using slight modifications to the protocol used by (Arumuganathan and Earle, 1991; Hermawaty *et al*., 2022). Two or three buds (∼50 mg FW) per biological replicate (three biological replicates) were dissected from the canes. The scales and woolly hairs surrounding the bud were carefully removed using forceps prior to chopping them with a razor blade in ice-cold nucleus-isolation buffer (NIB) at pH 7.4 (10 mM MgSO_4_, 50 mM KCl, 5 mM HEPES-K^+^, 6.5 mM DTT, 0.5 % (v/v) TritonX-100 and 3 % (w/v) PVP-40) on ice, and incubated on ice for 2 h with gentle swirling every half an hour. Following incubation, the suspension was passed through a 100 µm and 41 µm nylon mesh filter (Millipore, Germany), respectively, centrifuged at 450×*g* for 8 minutes at 4 °C. The supernatant was carefully discarded. The nuclei were resuspended in 2 mL NIB and treated with 50 µg.mL^-1^ of RNase A (Thermo Scientific, Australia, EN0531) for 6 min at room temperature before being stained with 20 µg.mL^−1^ propidium iodide and stored on ice until analysis. The ploidy of the nuclei was determined by fluorescence-activated sorting on a FACSCaliber flow cytometer (BD Sciences; Franklin Lakes, NJ, USA; https://www.bdbiosciences.com) at the Centre for Microscopy Characterisation and Analysis (University of Western Australia, Perth, WA). The flow cytometer was equipped with a primary blue 488 nm laser and data for ∼100,000 nuclei were recorded (i.e. until 20,000 G2 (4C) nuclei were collected). The proportion of nuclei with 2C and 4C DNA content was recorded. The data were visualised in real-time using scatter dot plots (forward scatter -FSC- and side scatter -SSC-) and histograms. The results were analysed using the Flowing Software (version 2, https://flowingsoftware.com) by manual gating to eliminate debris from the population of interest on the scatter plots and subsequent generation of histograms from the scatter plot data in which the different populations were gated to obtain the final proportion of G1, S and G2 nuclei computed by the software.

### Metabolite extraction and measurements

For each sampling date, two buds from nodes 6 and 8 were excised in the field and immediately collected in microtubes and placed on liquid nitrogen to be used for metabolite analysis (n=5). The frozen buds were ground to a fine powder at liquid nitrogen temperature. Water-soluble metabolites were extracted with chloroform-methanol as described previously (Lunn *et al*., 2006). Sugars were separated by hydrophobic interaction liquid chromatography HPLC using a Luna Omega Sugar column (3 µm, 100Å, 150 x 2.1 mm; Phenomenex, Torrance, CA, USA, www.phenomenex.com) fitted with a SUGAR (4 x 2.0 mm) security guard cartridge (Phenomenex) according to the manufacturer’s instructions, and quantified by tandem mass spectrometry on an AB Sciex 5500 Q-Trap triple quadrupole mass spectrometer (Feil and Lunn, 2018). Tre6P, other phosphorylated intermediates and organic acids were measured by anion-exchange high-performance liquid chromatography coupled to tandem mass spectrometry as described in Lunn et al. (2006) with modifications as described in Figueroa and Lunn (2016a).

### Statistical analysis

Analysis of variance was performed with data from at least four replicates in all cases and the means were compared using Tukey’s post-hoc honest significant difference test at the *P* < 0.05 level.

## Results

### Depth of dormancy through the year and the effect of weather conditions

We evaluated the depth of dormancy in buds from the southern hemisphere summer to spring (seven time points: January, February, April, May, June, August, and September). Hydrogen cyanamide (HC) treatment was used as a positive control for bud viability, as well as a reference for the depth of dormancy. The results showed that the depth of dormancy was greatest in late summer (February), with untreated buds taking 297 days to reach 50 % bud burst (BB50; Fig. 1A). Overall, from mid-summer to early winter the depth of dormancy was considerable, taking about 60 days or more to reach BB50. By June, the BB50 untreated buds burst within 25 days and the difference between the untreated and HC-treated positive control buds was negligible (Fig. 1A). January and February were the hottest months, with temperature maxima up to 39°C, and the coolest months were July and August with maxima below 20°C and minima as low as 4°C (Fig. 1B). Rainfall occurred more frequently from May to October, with occasional rainfall more than 40 mm per day in June and July (Fig. 1B). The mean temperature decreased from January to August, with a more significant decline occurring from April to June (Fig. 1C). During the period of greatest dormancy, mean temperatures were above 20°C (Fig. 1C). Environmental chilling, determined as PCU, did not accumulate appreciably until May, by which time the depth of dormancy had declined substantially from the peak in late February, and the difference between untreated and HC-treated buds was marginal (Fig. 1B). In this sense, we identified a period of physiological dormancy (commonly referred to as endo-dormancy) between February and March, and two periods of quiescence, one prior to dormancy (January) and the other after dormancy (from April), often called by many authors as para-dormancy and eco-dormancy. The initial decrease in BB50 from February to March did not correlate with climate data, although it is noted that the critical photoperiod for growth arrest is not well-defined in *V. vinifera*. The second decrease in the depth of dormancy (April to July) coincided with the decrease of mean temperatures and an increase in cumulative PCU (Fig. 1C). The average rainfall was low during the period of deepest dormancy (February) and only increased appreciably from April until July, declining thereafter (Fig. 1D). In this sense, we did not observe a clear correlation between the depth of dormancy and rainfall.

**Figure 1.**
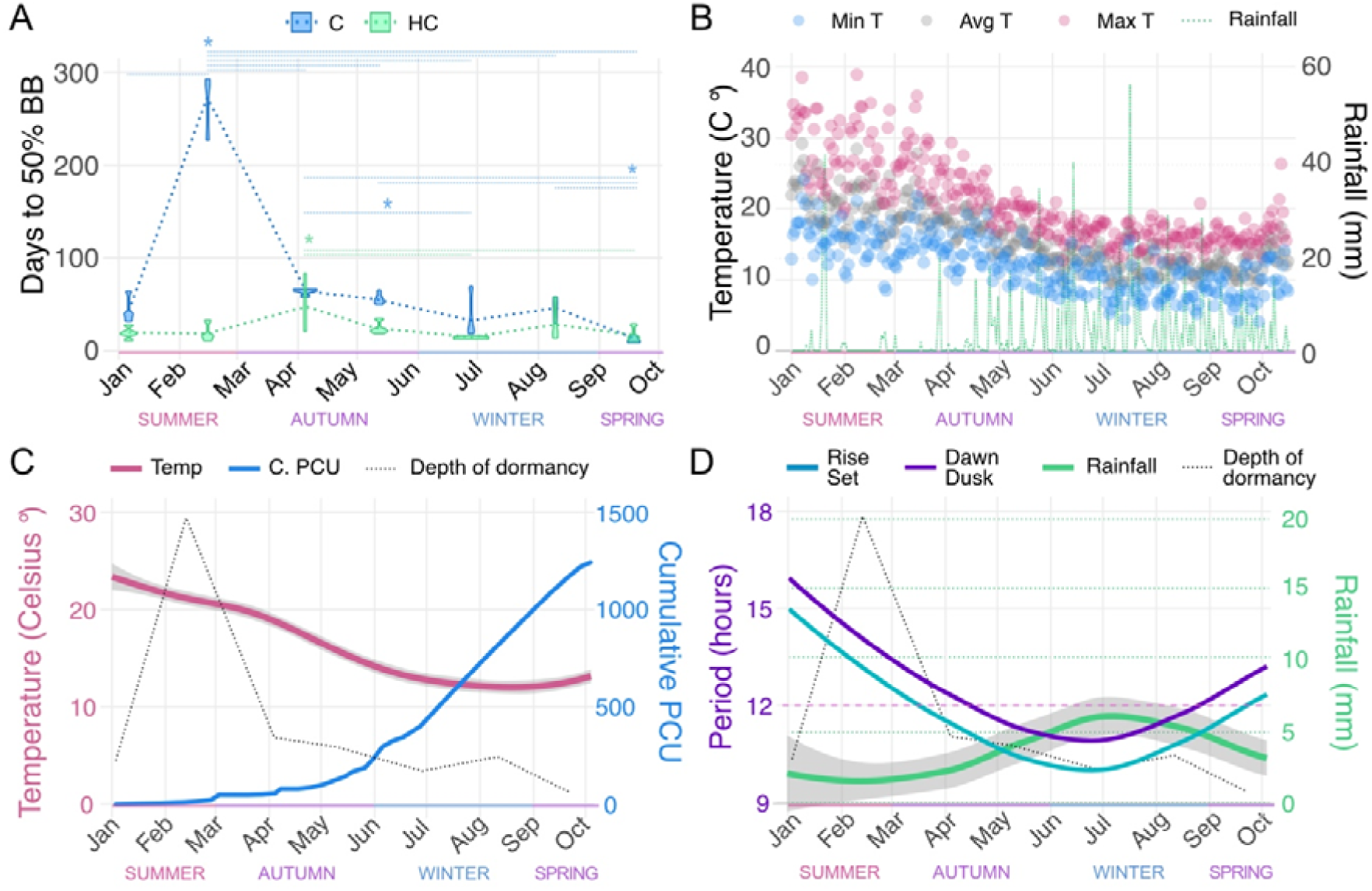
*Seasonal changes in perennating bud dormancy in grapevine*. **A.** Dynamic changes in the depth of dormancy of grapevine buds, grown as explants in permissive conditions and expressed as the time to 50 % bud burst (BB). Grapevine canes were taken at intervals during the southern hemisphere summer to spring (5 January, 15 February, 4 April, 10 May, 28 June, 9 August and 20 September). Single-node cuttings were treated with water (C) or 1.25% hydrogen cyanamide (HC) for 30 seconds and planted on a potting mix and subjected to forcing conditions in controlled growth chambers. The violins are the result of 5 replicates. Asterisks indicate statistical differences based on one-way analysis of variance with Tukey’s post-hoc honest significant difference testing at *P* < 0.05 for the samples connected with a light dashed line. **B.** Daily mean, maximum and lower air temperature represented by scatterplot and daily rainfall represented by a dashed line in Margaret River region of Western Australia in 2016. **C.** Relationship between the depth of dormancy, temperature annual trend and Positive Chill Units (PCU) accumulated. The dotted line represents the depth of dormancy as determined in panel A. Temperature is represented as a regression of the daily average temperature. The grey area represents +/- 95% confidence level intervals for prediction for the adjusted regression. **D.** Relationship between the depth of dormancy, rainfall annual trend and length of photoperiod. The dotted line represents the depth of dormancy as determined in panel A. Rainfall is represented as a regression of the daily rainfall. The grey area represents +/- 95% confidence level intervals for prediction for the adjusted regression. Photoperiod is represented as hours to sunrise/sunset and dawn/dusk. The horizontal dashed line at 12 h is plotted as a reference to see when days become shorter than nights. Meteorological seasons for the southern hemisphere temperate zone are marked in each panel.

### Moisture content and vascular connections

The moisture content and apoplastic connectivity of the buds were evaluated through the year. Figure 2A shows that the moisture content of the buds was higher in summer and then became lower from late summer to late winter, rising by spring (September). This behaviour is counterposed to the annual trend in rainfall (Figure 1D). The apoplastic path is vital to water transport in plants. As observed in Figure 2B, the buds were apoplastically isolated from the cane throughout this entire period, explaining the absence of correlation between rainfall and water content in the buds. It is noteworthy that this limited apoplastic transport is sustained until spring when some buds are already swelling (see September in Fig. 2B).

**Figure 2.**
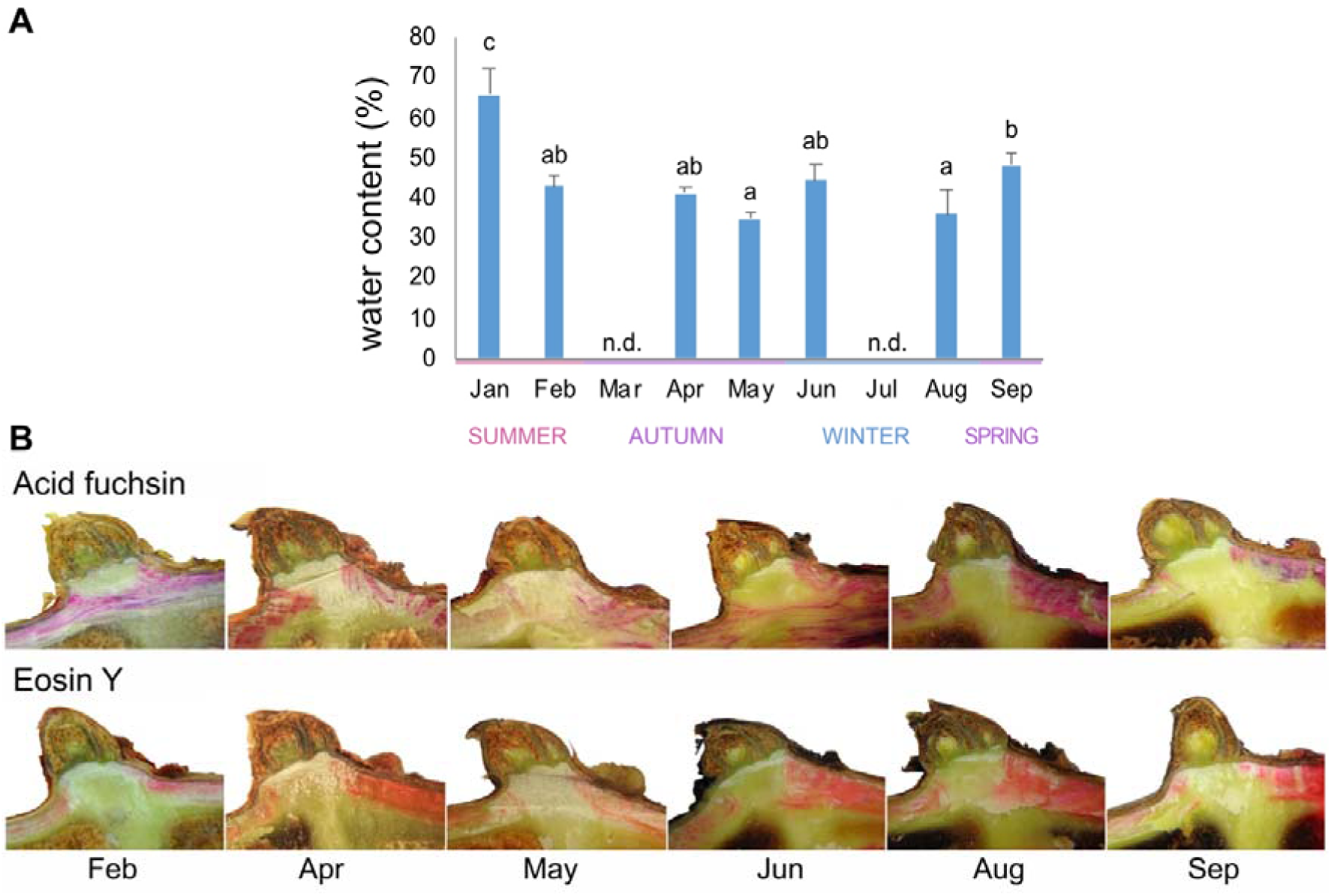
*Apoplastic isolation of buds throughout the season.* **A**. Moisture content. Bars indicate the mean of five biological replicates consisted of two buds each, error bars represent the standard deviation, and different letters show statistical differences based on one-way analysis of variance with Tukey’s post-hoc honest significant difference testing at *P* < 0.05, n.d., not determined; **B**. Apoplastic connectivity, as visualised by the acrotonic passage of two apoplastic dyes - acid fuchsin (upper row) and eosin Y (lower row).

### Internal oxygen profiles (pO_2_) and metabolic activity

Internal oxygen profiles (pO_2_) of the buds were also evaluated, as oxygen levels were suggested to regulate dormancy and bud burst (Meitha *et al*., 2015; Considine *et al*., 2017). In January and February, the internal oxygen partial pressure (pO_2_) was, on average, above 100 µM for the peripheral 1000 µm and then decreased to 25-50 µM in the inner part of the bud. During April, May and June, the levels of internal pO_2_ were more constant throughout the bud (∼100 µM), with a more pronounced decrease from 1500 µm in June. In September, however, the oxygen levels were greater for the outer sections of the bud than the inner section, at around 1500-2000 µm, where it was more hypoxic (Fig. 3A). The spatial variation within replicates is expected, considering the heterogeneity of the tissue, which comprises layers of bud scales alternated with trichomes (Signorelli et al., 2020).

**Figure 3.**
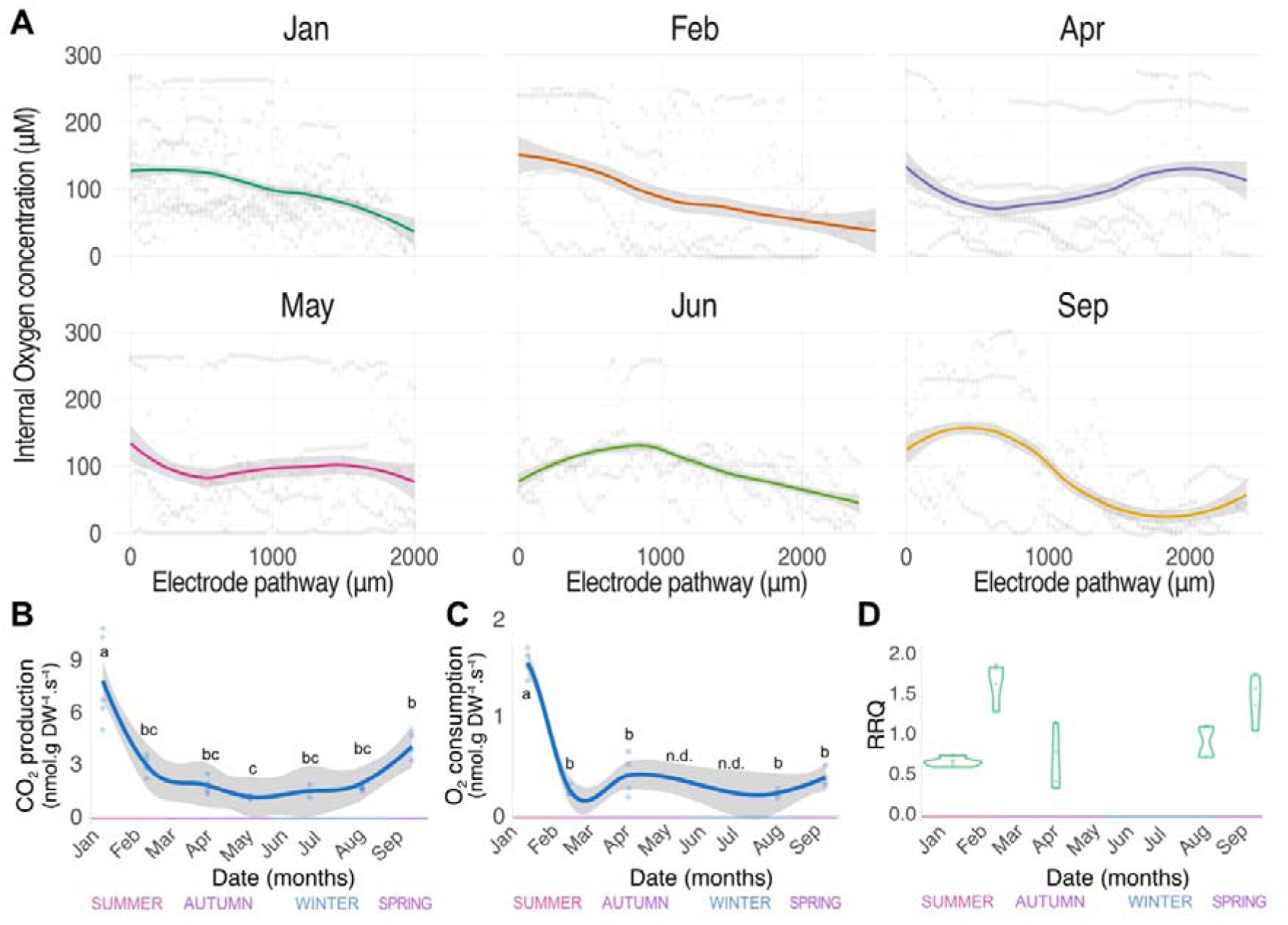
*Internal O_2_ concentration and respiration of buds along the year.* A, Internal oxygen profiles (pO_2_) represented as scatterplots of raw data (n > 4), with a regression curve applied (in colour) and 99 % confidence intervals (in grey shading); B, CO_2_ production (> 4); C, O_2_ consumption (> 4, except August n=2); D, relative respiratory quotient (RRQ). For A, B, and C, individual observations are plotted as light dots, the line is an adjusted regression of the data, and the grey regression represents +/- 95% confidence level intervals for prediction for the adjusted regression. Different letters show statistical differences based on one-way analysis of variance with Tukey’s post-hoc honest significant difference testing at *P* < 0.05. For D, the data is represented as a violin plot, with individual observations plotted as light green dots.

The metabolic activity was followed by measuring the CO_2_ release and O_2_ consumption (Fig. 3B-C). The highest metabolic activity was found in summer (January) before both CO_2_ production and O_2_ consumption declined. For both parameters there was an increase from May to September, however, this was only statistically significant for CO_2_ release. The greater increase in CO_2_ release relative to O_2_ consumption towards the end of the season results in a slight increase of the relative respiratory quotient (RRQ, Fig. 3D), which may indicate a change towards metabolism of more oxidised substrates or use of reducing power for alternative processes. Likewise, the low O_2_ consumption found during the dormancy peak (February), with a CO_2_ release that was moderate, resulted in a high RRQ (Fig. 3D).

### Cell cycle

The population of cells in the different phases of the cell cycle was evaluated along the year to determine whether there was any correlation between the quiescent state of the bud and the cell cycle (Fig. 4). In January about 54% of cells were in the G1 phase, 37% in S phase and 8% in G2 phase. In the transition to late summer (January-February), when the depth of dormancy was greatest, the number of cells in the S phase significantly declined. After this period, the fraction of cells in S phase remained unchanged until bud burst, at about 30%. The fraction of cells in G1 remained unchanged from January to late June with values about 50%, in line with a number of studies showing that the cell cycle arrests in the G1 phase in dormant tissues (Velappan et al., 2017). However, in August and September the fraction of cells in G1 had declined to about 38%, coupled with an increase in the fraction of G2 cells. This change in the distribution of cells at each phase of the cell cycle suggests that dormancy release had been initiated. The G2/G1 ratio is a common metric of cell division. Here we observed that August and September saw a two-fold increase in the G2/G1 ratio, indicating a significant shift towards proliferation (Fig. 4B). Notwithstanding, these data have no spatial resolution among the heterogenous tissues that comprise a primordial shoot.

**Figure 4.**
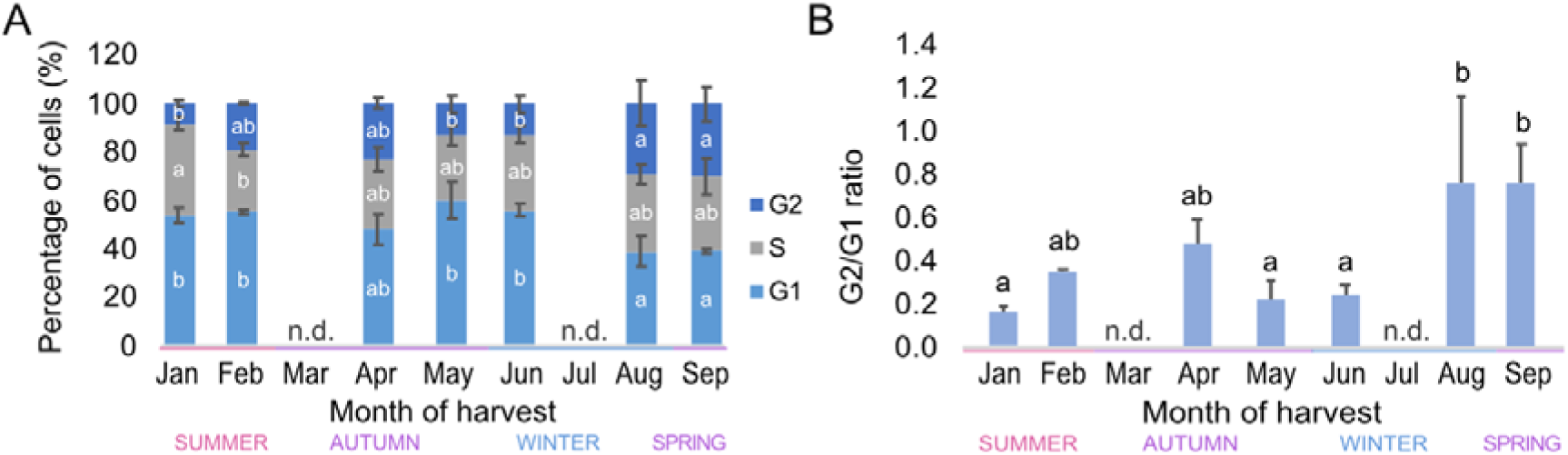
*Variation of the cell cycle phases through the year* A. Cell cycle phases per month. B. G2/G1 ratio. Bars indicate the mean of five biological replicates consisted of two buds each, error bars represent the standard deviation, and different letters show statistical differences based on one-way analysis of variance with Tukey’s post-hoc honest significant difference testing at *P* < 0.05, n = 3.

### Analysis of sugars and metabolic intermediates during the seasonal changes

We investigated seasonal changes in the levels of sugars and respiratory intermediates in the buds. A principal component analysis (PCA) of our metabolite data showed that January and February are clearly separated from the other months, while all the other months clustered together with a few replicates being away from their group (Fig. 5A). In this sense, the PCA of all metabolite data distinguished the samples by depth of dormancy, as February and January were the months where the deepest dormancy was found (Fig. 1). We performed a hierarchical cluster analysis of the metabolite data for all the months and replicates. In general, we observed that most sugars with an energetic function, sugar-phosphates, and tricarboxylic acids peak in January, their levels are low during the period of quiescence until September when many of them increase again (Fig. 5B). Whereas the sugars that have been associated with freezing protection and osmotolerance had lower concentration during the summer (January-February) and spring (September) periods and greater concentrations during the autumn and winter period (April to August), consistent with their functions and the lower water content (Fig. 2A, 5B). As expected, myo-inositol, which is used to produce galactinol and raffinose-family oligosaccharides has a contrasting pattern.

**Figure 5.**
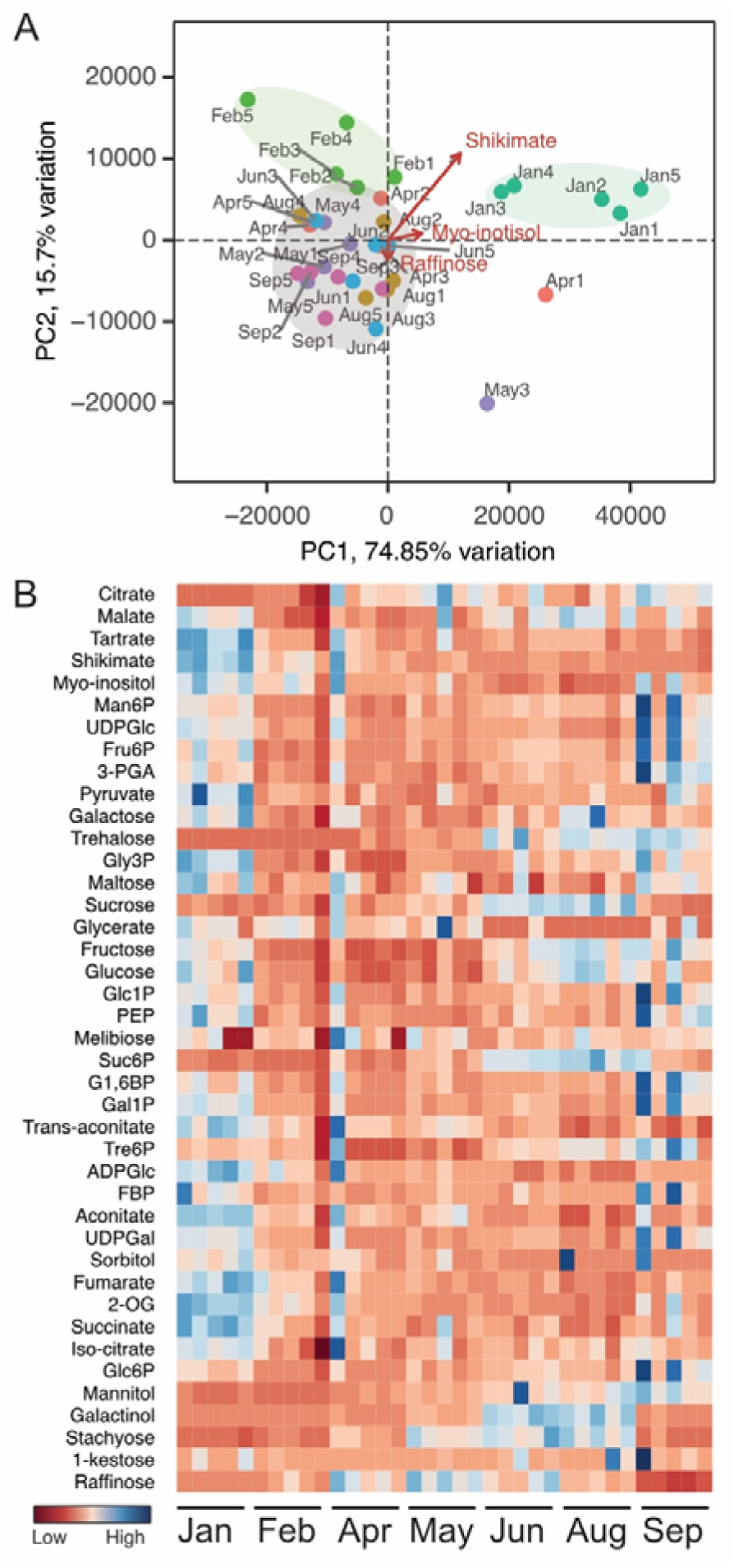
*Principal component analysis (PCA) and hierarchical cluster heatmap of metabolites from grapevine buds along the year.* A. PCA using all metabolite data. Samples from different months are represented in different colour and labelled accordingly. Ellipses highlight either replicates from the same month that cluster together and separate from other months (e.g. Jan and Feb), or samples from different months that group closely in the PCA space. The eigenvectors in magenta show the contribution of the main variable (metabolite) driving the principal component. B. Heatmap, the values were scaled per each metabolite, and the order of the months was fixed. Five replicates of grapevine buds were used to determine the concentration of metabolites.

When only sugars were considered for the PCA, we observed that not only January and February were clearly distinct to the other samples but also September (Fig. 6A). Given that many of these sugars have an energetic function, it is not surprising that these months are separated from the others given that we observed higher humidity and metabolic activity (Fig. 2A and 2B).

**Figure 6.**
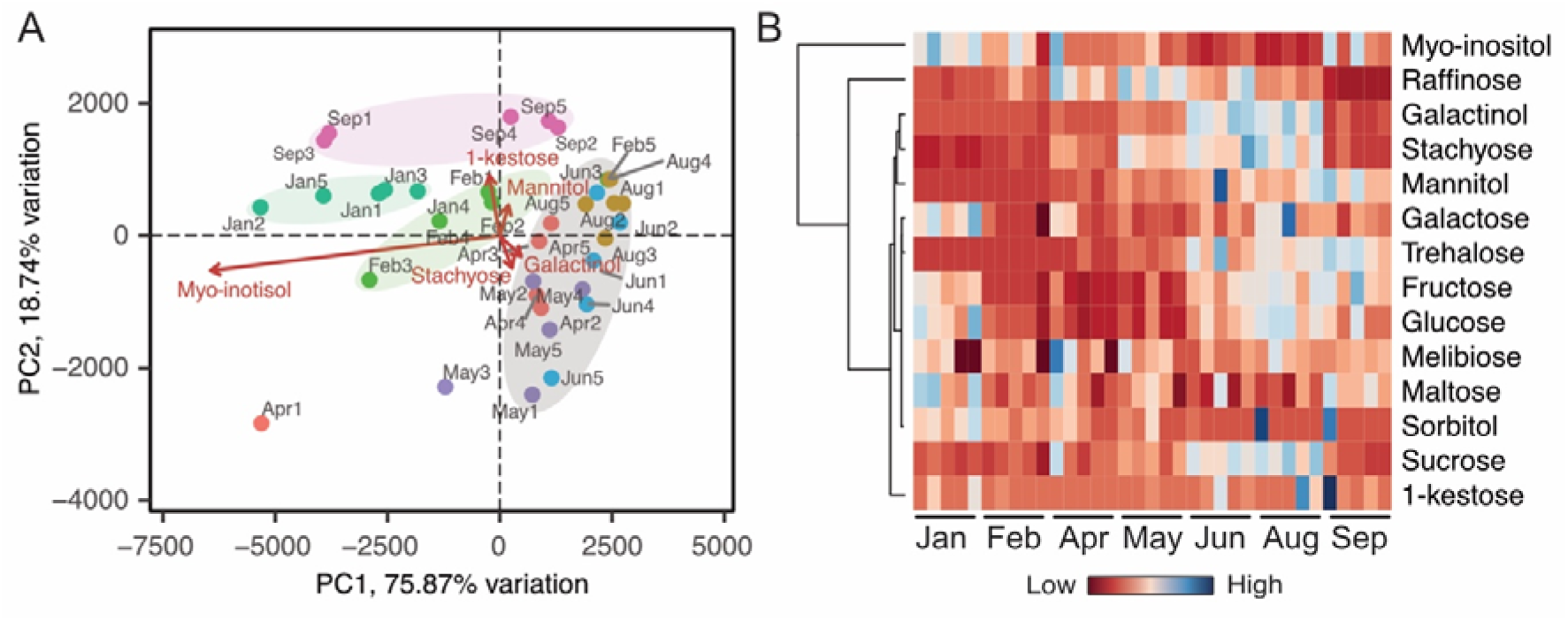
*Principal component analysis and hierarchical cluster heatmap of sugars along the year.* A. PCA using only sugars data. Samples from different months are represented in different colour and labelled accordingly. Ellipses highlight either replicates from the same month that cluster together and separate from other months (e.g. Jan, Feb and Sep), or samples from different months that group closely in the PCA space. The eigenvectors in magenta show the contribution of the main variable (metabolite) driving the principal component. B. Heatmap, the values were scaled per each metabolite, and the order of the months was fixed.

The hierarchical cluster analysis of these metabolites showed myo-inositol did not group with any of the other metabolites (Fig. 6B). Raffinose was also separated from the other sugars, and then all the other sugars clustered together, with 1-kestose displaying a more distinct profile (Fig. 6B).

To further understand the profile of the metabolites and their absolute value, we plotted each metabolite per month, grouped as sugars, sugar alcohols, and hexose-phosphates (Fig. 7), and glycolytic and tricarboxylic (TCA) cycle intermediates (Fig. 8). Looking at their concentration, sucrose, tartaric acid, citrate, glucose and fructose were the most abundant metabolites in the buds (Fig 7 and 8). Glucose and fructose followed a very similar pattern characterized by high values in January and August-September (Fig. 7), when respiration rates were also high (Fig. 3B-C), however, the ratio between them was not always constant, as we observed a significant decrease of the [Glc]/[Fru] ratio towards the budburst period (Fig. 9). Galactose showed the same profile as glucose and fructose but at lower concentrations (Fig 7). Sucrose and its phosphorylated form (Suc6P) gradually accumulated from low levels in January to a maximum in July/ August, which preceded the peak of glucose and fructose. The results suggest that the buds accumulate sucrose through winter, and prior to bud burst sucrose is catabolised into glucose and fructose which are further catabolised to provide energy.

**Figure 7.**
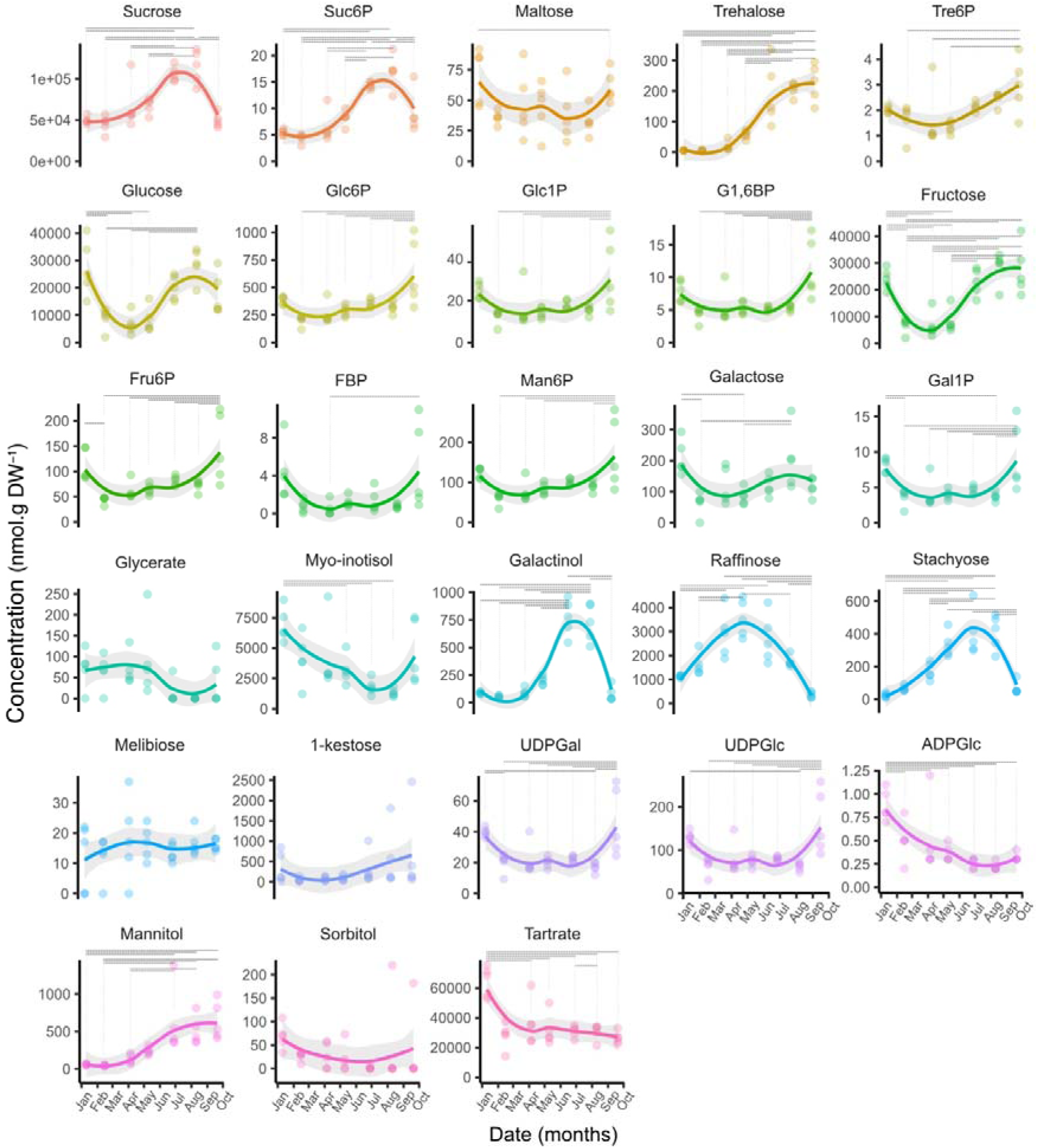
*Concentration of sugar, sugar alcohols and hexose-phosphates in grapevine buds at different stages of the year.* The horizontal dotted bars connecting different months indicate statistical differences based on one-way analysis of variance with Tukey’s post-hoc honest significant difference testing at *P* < 0.05, n=5. The grey area represents +/- 95% confidence level intervals for prediction for the adjusted regression.

**Figure 8.**
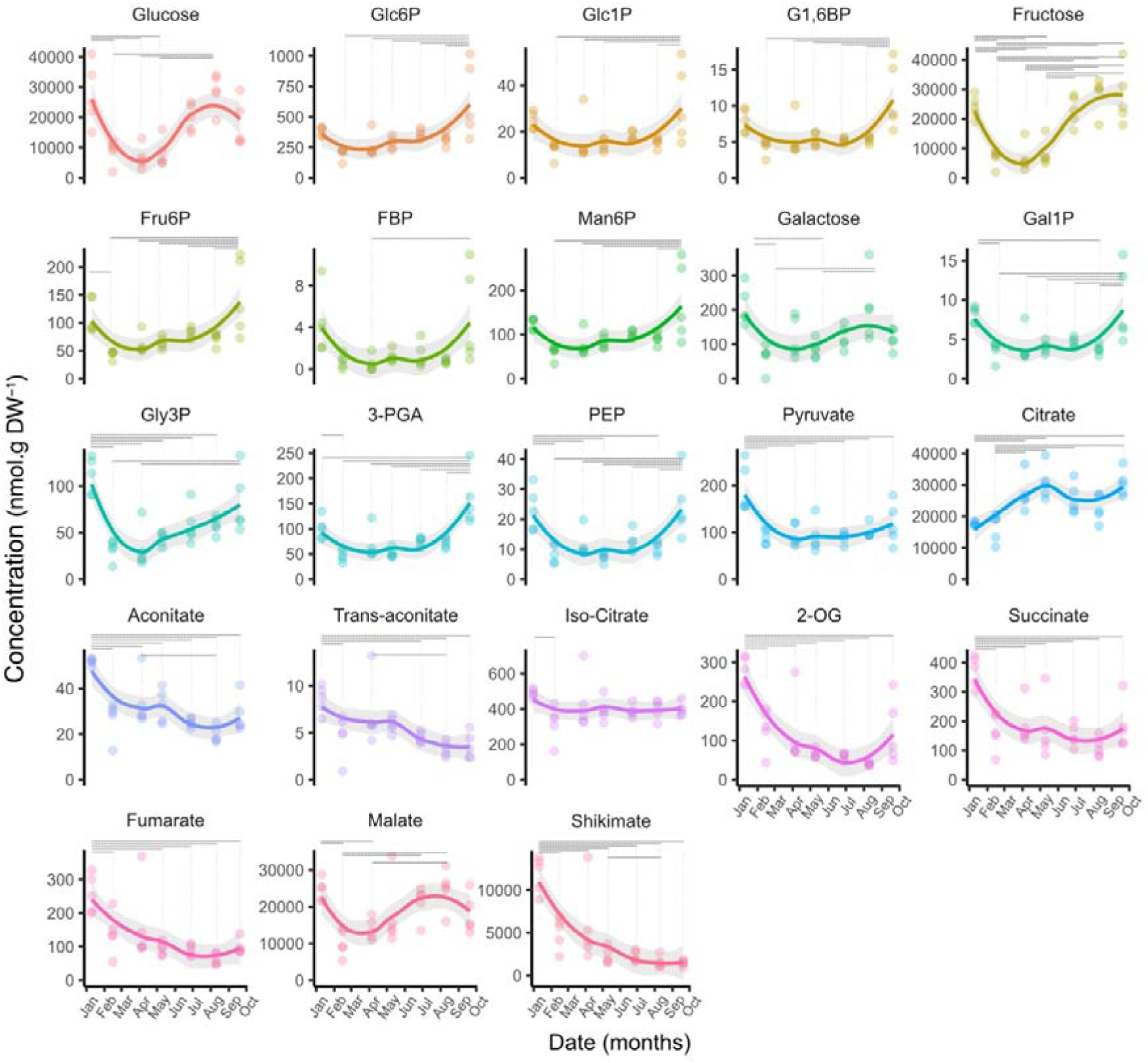
*Concentration of glycolytic and TCA-cycle intermediates in grapevine buds at different stages of the year.* The horizontal dotted bars connecting different months indicate statistical differences based on one-way analysis of variance with Tukey’s post-hoc honest significant difference testing at *P* < 0.05, n=5. The grey area represents +/- 95% confidence level intervals for prediction for the adjusted regression. The hexoses and hexose-phosphate data are the same as those reported in Fig. 7, and are reproduced here to allow direct comparison with downstream glycolytic intermediates.

**Figure 9.**
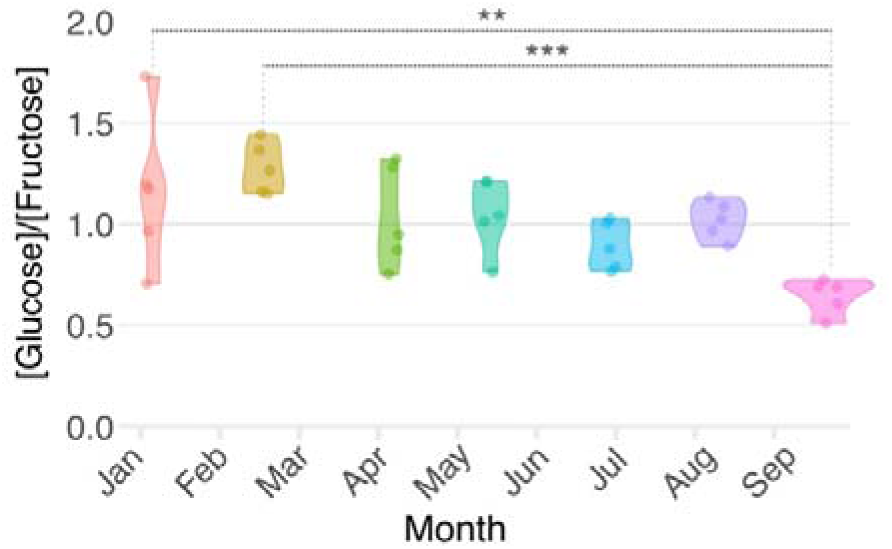
*Glucose to fructose ratio in grapevine buds throughout the season.* The violin plots show data from five replicates (represented in circles). The horizontal dotted bars in black connecting different months indicate statistical differences based on one-way analysis of variance with Tukey’s post-hoc honest significant difference testing. The asterisks indicate statistical differences based on one-way analysis of variance with Tukey’s post-hoc honest significant difference testing at *P* < 0.01 (**) or *P* < 0.01 (***), n=5.

The accumulation of monosaccharides in spring is key to promote bud burst, as they will act as energy substrates to sustain growth but also the conversion of sucrose to glucose and fructose will double the osmotic potential, thereby increasing the turgor pressure and driving cell expansion. Moreover, these sugars and some of the phosphorylated intermediates of sugar metabolism may play a role as signalling molecules to promote bud outgrowth. Supporting this idea, we observed a significant increase of all phosphorylated monosaccharides (Gal1P, Glc6P, Glc1P, G1,6P, Fru6P, Fru1,6P, Man6P) as well as trehalose 6-phosphate (Tre6P) in September (Fig. 7). In general, most glycolytic intermediates showed a similar profile (Fig. 8), being statistically higher either in January, September or both months, which is consistent with the greater respiration rates we observed in these months.

The TCA cycle intermediates displayed a greater concentration in January than the rest of the year, except for citrate which had lower levels during the summer period (Fig. 8). This suggests that the TCA cycle is most active in summer but remains less active during the rest of the year. Malate displayed a different profile, showing the lowest levels in February and April, when the depth of dormancy is greater.

Sorbitol and mannitol showed a similar pattern to trehalose, progressively increasing from April/May (Fig. 7). Another group of molecules showing a similar pattern were galactinol, raffinose and stachyose (Fig. 7). Galactinol is a precursor of raffinose, which in turn is the precursor of stachyose, so the similar pattern is not surprising, however, it indicates a flow of these sugars all the way through to stachyose or more complex oligosaccharides (e.g. verbascose) that we did not measure. In addition, myo-inositol, a precursor of galactinol, show an opposite pattern to galactinol (Fig. 7), suggesting a possible use of myo-inositol through the year to produce galactinol until July/August, when galactinol levels are the highest and myo-inositol levels the lowest, and then galactinol is catabolised and its biosynthesis reduced resulting in an increase of myo-inositol prior to bud burst (Fig. 7). Similarly to myo-inositol, shikimate and tartrate, two abundant metabolites in the buds, progressively declined from their peak abundance in January (Fig. 7). No significant change in concentrations were observed in melibiose and 1-kestose throughout the experimental period (Fig. 7). UDP-galactose and UDP-glucose shared a common profile, with concentration peaks in January and September, whereas ADP-glucose declined continuously from its peak in January (Fig. 7). ADP-glucose is essential for starch biosynthesis. During late autumn and winter, the absence of leaves does not allow the plant to produce sugars for the build-up of starch reserves, so it is not surprising the levels of ADP-glucose decrease. On the other hand, UDP-Glc levels significantly increased prior to the bud burst period. Additionally, UDP-Glc is known to be produced from sucrose degradation in sink tissues to build up cell wall synthesis (Wai *et al*., 2017). UDP-Glc has also been suggested to play a signalling role during development (Janse van Rensburg and Van den Ende, 2018).

## Discussion

### The physiological and biochemical data delineate three periods of quiescence in grapevine buds

Here we analysed physiological features of *V. vinifera* cv. Cabernet Sauvignon buds over a seven-month period of quiescence, from the onset of dormancy to immediately before bud burst. Late summer (January to February) was characterized by the establishment of the peak of dormancy. At this stage, the basal buds are lignified but the internal process of dormancy was only beginning to establish, as evident in the increase in time to bud burst over this period. The literature commonly refers to this period as the transition between para-dormancy and endo-dormancy. This peak of dormancy was characterized by an apoplastic isolation of the bud from the cane, a reduction of the moisture content of the bud, a decrease in the proportion of cells in the S phase of the cell cycle, and a reduction of the respiration rate (CO_2_ release and O_2_ consumption). High levels of myo-inositol, shikimate, tartrate, ADP-Glc, and some TCA cycle intermediates were signature features of this first period of quiescence (Fig. 5, 7 and 8). Conversely, the levels of citrate were the lowest during this initial period of quiescence (Fig. 8).

In the second major period of quiescence, the depth of dormancy declined in a biphasic manner; it rapidly declined between the end of summer and mid-autumn, followed by a more moderate decline towards winter. These dynamics cannot be explained by the chilling or rainfall conditions, as chilling only began from mid-late autumn and rainfall also became more frequent after that period. However, daylength was already noticeably shorter than in summer (Fig. 1D). In hybrid aspen, short photoperiods were shown to increase ABA levels and expression of ABA receptors, leading to the induction of dormancy, the closure of plasmodesmata and prevention of growth-promoting signalling to reach the meristem (Tylewicz *et al*., 2018). Here, respiration, internal pO_2_, and moisture content of buds declined progressively from summer to a point of inflection in early winter. These patterns are consistent with an earlier study in cv. Merlot in the same vineyard (Velappan *et al*., 2022b). During this second period of quiescence, most metabolites remained unvaried; however, high levels of raffinose were a clear metabolic signature of this period, as well as a steady increase in stachyose, sucrose, sucrose-6P, galactinol and trehalose (Fig. 7).

The apoplastic pore size subtending the buds, which was determined to exclude molecules greater than 2.1 nm (Signorelli *et al*., 2020b), was restricted throughout the study period, even when buds had begun to swell in spring, while the most pronounced shift in cell division was the increased proportion of G2 cells in late winter, relatively coincident with bud moisture content. This final transition accompanied a considerable increase in the ratio of G2/G1 cells, which is a common proxy for the mitotic index (Velappan *et al*., 2022d). We observed peaks in metabolic activity in January and September. However, the G2/G1 ratio suggests that cells in January are likely still growing but not dividing (low G2/G1 ratio, Fig. 4), while in September cell division has been initiated (high G2/G1 ratio, Fig. 4). In addition, during this third phase, the depth of dormancy was reduced to the minimum, the moisture of the bud increased and so did the respiration rate.

The internal pO_2_ fluctuated during these three periods, being reasonably uniform through the bud profile in the second period (February to June), whereas in the first and latest periods, summer (January) and early spring (September), the level of oxygen declined to below 20 µM in the core of the bud (Fig. 2C). This could be attributed to a higher metabolic activity of the bud in those two months, which we confirmed by respiration measurements (Fig. 3B-C), which increases O_2_ demand and leads to a decrease in internal O_2_. Based on our previous work suggesting that internal tissues are not a significant barrier to O_2_ diffusion (Signorelli et al. 2020), we consider respiratory demand the primary driver of this hypoxia. On the other hand, the greater oxygenation of the bud in the outer area in September could be linked to the fact that the outer scale and trichomes become looser prior to bud burst, allowing ready influx of O_2_. In terms of metabolites, towards the end of this phase peak levels of trehalose, 3PGA, UDP-Gal, UDP-Glc, and many sugar phosphates such as Tre6P, Glc6P, Glc1P, G1,6BP, Fru6P, Man6P were found. All in all, these data broaden our understanding of bud quiescence or the so-called endo-, para-, and eco-dormancy periods from a physiological and biochemical view.

### The dynamics of metabolites in grapevine buds through the season

In annual herbaceous plants, such as arabidopsis and sorghum, the role of sugars and some sugar-phosphates in dormancy and bud outgrowth control is well established (Barbier et al., 2015; Fichtner et al., 2017). In general, low sugar content has been demonstrated to inhibit bud outgrowth (Mason *et al*., 2014; Kebrom and Mullet, 2015). However, little is known about the endogenous levels of sugars within grapevine buds and their dynamics across the season. Despite the buds being quiescent, the concentrations of sugars, sugar phosphates, TCA cycle intermediates, and other metabolites varied significantly during that period. Based on our PCAs, we could separate our samples into up to four subgroups (Fig. 6), January, February, April-August, and September, so we defined these groups as four different metabolic phases to organize the discussion of the metabolite data.

### Several monosaccharides, sugar phosphates and most TCA cycle intermediates decrease during the onset of the quiescent period in grapevine buds

The first metabolic phase, in early summer, was characterised by the highest concentrations for ADP-Glc, tartrate, most TCA cycle intermediates, pyruvate, myo-inositol and shikimate (Fig. 7 and Fig. 8). Maltose, glucose, fructose and galactose, as well as their phosphorylated forms also had high values during this phase (Fig. 7). Physiologically, this phase represented the onset of dormancy (Fig. 10). The decrease of hexoses, with high levels of pyruvate, and TCA cycle intermediates, together with the higher respiration rate during January points to metabolism of the buds being highly active in early summer. Tartrate, which is known to reach high levels in grape berries, also reached high concentrations in the bud during summer. Radiolabelled tartrate is recovered as carbon dioxide released by berries, indicating that tartrate is catabolized in grapevines (Hrazdina *et al*., 1984). Thus, tartrate is likely being catabolized as a source of carbon and energy to support metabolic activity in the developing bud. Shikimate is a precursor for phenylpropanoids (e.g., flavonoids, isoflavonoids, coumarins, aurones, stilbenes, catechin, lignin), many of which have protective functions against abiotic stress, deterrence of insect pests and pathogen defence. The very high initial shikimate content and decrease over the seasons (Fig. 7) suggests a substantial investment in defence compounds to protect the bud throughout the rest period. Similarly, chloroplasts within the leaf primordium are likely accumulating starch as a reserve for the rest of the season as suggested by the high levels of ADP-Glc (Fig. 8). Starch is known to have high concentrations in the bud which rapidly decrease during the bud burst period (Meitha *et al*., 2018). Likewise, myo-inositol is needed for the biosynthesis of complex sugars, such as raffinose, which were accumulated in the following months. Myo-inositol and its derivatives are also considered crucial compounds for development and signalling in plants (Valluru and Van den Ende, 2011; Roitman and Eshel, 2024); therefore, a signalling role in the transition between para-dormancy and endo-dormancy or during bud burst should not be ruled out.

**Figure 10.**
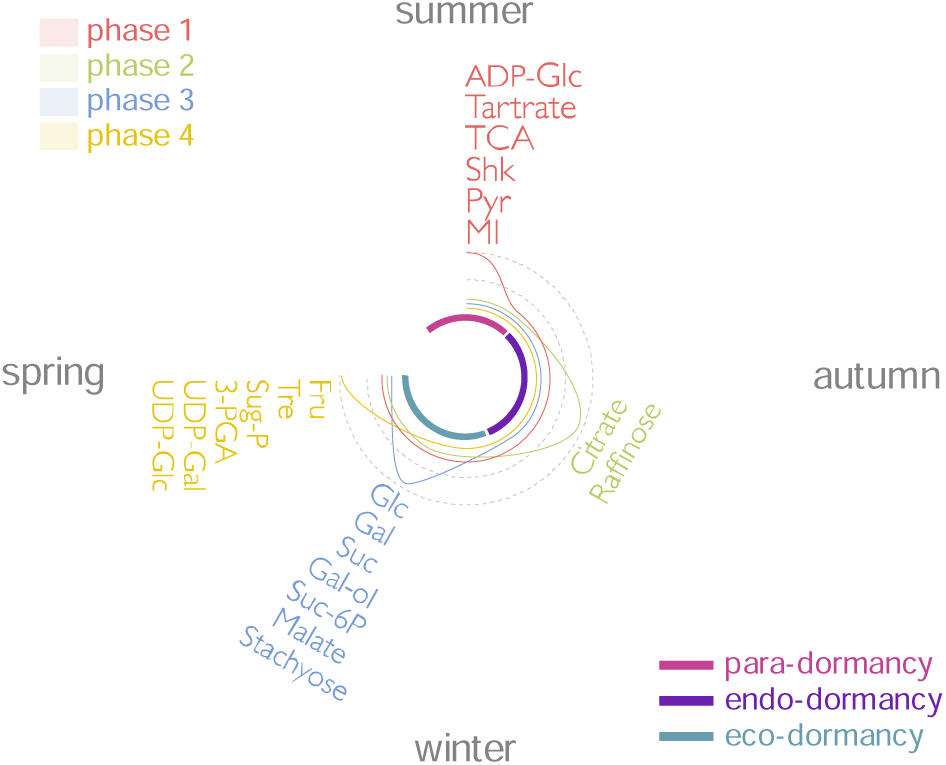
Schematic representation of metabolite profiles along the season and the correlation of their peak concentrations with the seasons and the quiescent state. The different metabolic phases were defined based on the PCA for the metabolomic data. 3-PGA, 3-phospho glycerate; Fru, fructose; Gal, galactose; Gal-ol, galactinol; Glc, glucose; MI, myo-inositol; Pyr, pyruvate; Shk, shikimate; Suc, sucrose; Sug-P, sugar-phosphates; TCA, tri-carboxylic acid cycle intermediates; Tre, trehalose.

### The peak of dormancy correlated with the lowest levels of sugars and sugar-phosphates in grapevine buds

The second metabolic phase (February or summer) coincided with the peak of dormancy (Fig. 1) and represented a period in which almost all metabolites remained unvaried except for citrate and raffinose, which were the only two metabolites that were increasing (Fig. 7, 8 and 10). Thus, both the metabolite data and the physiological dormancy suggested the deepest rest during this month. The levels of many sugars, such as glucose, fructose, trehalose, and sucrose, were at the minimum level during this second phase. In concordance, gene ontologies representative of the transcriptional transition from summer to autumn indicated a strong signature of response to metabolic stimuli, including starvation and nutrient levels (Velappan *et al*., 2022b).

### Raffinose and citrate accumulate at the begin of the period of cold, while several sugars and sugar alcohol accumulates towards the end of this period in grapevine buds

The third metabolic phase was the longest one, from late summer to late winter, and was characterized by the peak of raffinose and citrate in autumn (May) and the subsequent accumulation of many metabolites whose concentrations peaked in late winter (August) (Fig. 10). The build-up of raffinose coincided with the period of physiological dormancy or endo-dormancy (Fig. 10), reaching its peak in late autumn. After that period, when the day length is the shortest and rainfall increases, raffinose levels start to decrease again (Fig. 1D and Fig. 7). Raffinose accumulation has been documented as a response to cold (Tarancón *et al*., 2017)and it has been linked to freezing tolerance in grapevine (De Rosa *et al*., 2022). In seeds, raffinose has been linked to seed longevity (Salvi *et al*., 2022). Therefore, raffinose could be accumulating in cv. Cabernet Sauvignon buds to protect the buds during the winter period. In cv. Cabernet Sauvignon leaves, a few hours of cold exposure produce a rapid decrease in raffinose concentration (Jiang *et al*., 2016a). We observed a continuous decrease from winter, suggesting that also in the cv. Cabernet Sauvignon buds raffinose is catabolised during the cooler period. In canes of the grapevine cv. Vinhão, a gene involved in raffinose biosynthesis (*VviRafS5*) was shown to be induced upon cold exposure (Noronha *et al*., 2022). This could either mean that this cultivar responds to cold accumulating raffinose or that raffinose is produced in the cane to feed the bud during the dormancy period where it is catabolized. In future research, it would be good to know in one cultivar the levels of raffinose in the cane and bud at the same time and the response of the transcripts involved in its metabolism. The response of raffinose metabolism was shown to be genotype dependent, as in ‘Beta’ (*Vitis riparia × V. labrusca*) leaves cold-induced raffinose levels (Jiang *et al*., 2016b). In a transcriptomic study, we observed that cv. Merlot buds displayed a strong decline in the expression of homologues of *RAFFINOSE SYNTHASE* in the transition from summer to autumn (Velappan *et al*., 2022b), suggesting either that raffinose biosynthesis is differentially regulated in buds of different cultivars or that post-translational regulation may prevail upon raffinose family oligosaccharide synthesis.

In winter (June-August), we found a higher content of sucrose, Suc6P, galactinol, stachyose, mannitol, and trehalose (Fig. 7), and this was immediately followed by an increase of glucose, galactose, fructose, and malate in late winter (Fig. 7, 8 and 10). Soluble sugars such as glucose, fructose, and sucrose are considered to be induced during the period of quiescence thanks to the activation of amylase activity by chilling which degrades starch reserves (Monteiro *et al*., 2022). Invertase and SWEET sugar transporters have been shown to be activated to increase sugar content in buds and promote bud outgrowth (Girault *et al*., 2010; Rabot *et al*., 2012, Kebrom and Mullet, 2016), and the induction of these enzymes was linked to light perception (Kebrom and Mullet, 2016). We observed that the accumulation of these sugars coincided with the increase in day length (Fig. 1D and 7). Therefore, it is possible that also in grapevine buds photoperiod contributes to regulating the expression of enzymes important to augment sugar content within the bud.

The concomitant accumulation of molecules such as stachyose and sucrose, accompanied by the decrease of raffinose suggest that during this phase increasing sucrose provides greater substrate availability for raffinose synthesis, although raffinose itself may not accumulate because it is mostly converted into stachyose or higher degree of polymerization raffinose-family oligosaccharides. Sorbitol, mannitol, and trehalose showed a similar pattern progressively rising from April/May (Fig. 6). These sugars are known to be compatible osmolytes with kosmotropic activity, and trehalose has a well-known role as an anti-freezing molecule (Garg *et al*., 2002), so it is possible that these molecules are accumulated through the year to protect the bud from freezing in winter. In addition, the buds showed the lowest levels of water content in autumn, so their accumulation can be also a response to osmotic stress.

Sucrose can promote growth both as a source of energy and as a signalling molecule (Barbier *et al*., 2015). Since we did not observe an activation of metabolic activity (Fig. 4) and the products of its catabolism (glucose and fructose) until the end of this phase three, we conclude that sucrose is accumulated but not catabolized until the end of this phase when energy is required for growth resumption.

Besides, the high levels of glucose, sucrose, and Suc6P reached prior to being catabolized could play a signalling role, for example, in activating genes essential for growth resumption via the bZIP11 transcription factor (Ma et al., 2011) and regulation of the SUCROSE-NON-FERMENTING1-RELATED PROTEIN KINASE 1 (SnRK1; (Tarancón *et al*., 2017). In arabidopsis root hair, glucose was shown to induce TOR activity which can phosphorylate E2Fa to promote root cell proliferation (Xiong *et al*., 2013b). Thus, the increase of glucose in the weeks prior to bud burst could be playing a similar mechanism in grapevine buds. Moreover, the increase of Tre6P prior to the bud burst period would inhibit SnRK1 (Baena-González and Lunn, 2020), reducing the inactivation of TOR through phosphorylation mediated by SnRK1, and further promoting the induction of metabolic activity by TOR. In this sense, the concomitant increase of Tre6P and decrease of sucrose that we observe prior to bud burst aligns with the current models suggesting that Tre6P can result in reduced levels of sucrose, either by altering sucrose allocation or reducing starch degradation (Göbel and Fichtner, 2023), and play a role in promoting bud outgrowth, models that were mainly originated from model plants such as arabidopsis or pea.

### Sugar-phosphates accumulate prior to bud burst in grapevine buds

The fourth and last metabolic phase was defined by the highest concentration of fructose, trehalose, most of the measured sugar-phosphates, 3-PGA, UDP-Gal, and UDP-Glc (Fig. 7 and 8). At this stage, it is likely that sucrose and glucose are being respired to fuel the bud outgrowth through glycolysis, as we observed an increase of most glycolytic intermediates studied here (Glc6P, Fru6P, FBP, 3-PGA, and PEP). This rise in glycolytic intermediates accompanies a rise in respiratory CO_2_ production (Fig. 4), which indicates there is increased flux through the TCA cycle and, by inference, increased glycolytic flux to feed carbon from sugars into the TCA cycle. The lower [Glc]/[Fru] ratio observed toward the bud burst period may suggest a preferential use of glucose, either to fuel metabolic activity or to support cell wall biosynthesis, as indicated by the concurrent increase in phosphorylated glucose (Fig. 8) and UDP-glucose levels (Fig. 7).

In an explant experiment we showed that light downregulates the expression of one of the *TREHALOSE-6-PHOSPHATE PHOSPHATASE* (*TPP*) genes (*VIT_01s0011g05960, Vitvi01g00509 in v4.3*) in explant grapevine buds (Signorelli *et al*., 2018). TPP converts Tre6P into trehalose. Here, we observed that the accumulation of trehalose levels stops with the increase of day length (from August to September) while an increase in UDP-Glc occurs, suggesting that the downregulation of TPP mediated by light also occurs in the field. The decrease in trehalose accumulation observed from August to September might also implicate the production of Tre6P, which we observed to reach the greatest levels in September. Tre6P is considered to play a signalling role that promotes growth of sink organs when sucrose supplies are abundant (Figueroa and Lunn, 2016b), in part through the inhibition of SnRK1 to promote growth (Schluepmann *et al*., 2003; Zhang *et al*., 2009; Delatte *et al*., 2011). In another transcriptome study of cv. Merlot, we did not observe major changes in genes related to trehalose metabolism until the transition from winter to spring (Velappan *et al*., 2022b).

### Possible metabolic markers to differentiate quiescent states in grapevine buds

We looked for metabolic signatures for the three different period of quiescence (i.e. quiescence prior to dormancy, or para-dormancy; physiological dormancy, or endo-dormancy; and quiescence after dormancy, or eco-dormancy) in grapevine buds of cv. Cabernet Sauvignon. We observed that levels of shikimate, tartaric acid, fumarate, succinate, aconitate, trans-aconitate, or 2-oxoglutarate were high only during the quiescence period prior to dormancy, whereas, sucrose, trehalose, galactinol, stachyose, and mannitol were exclusively high during the period of eco-dormancy. Low levels of hexoses (glucose, fructose, galactose) and an increase in raffinose concentrations were features of the endo-dormancy period. If this is extrapolated to other grapevine cultivars, the sugar profiles of buds could be used as a proxy to differentiate the period of dormancy (endo-dormancy) and quiescent states (para- and eco-dormancy). Such readily measured markers for the different dormancy states could be of great practical value to researchers studying woody perennials.

## Conclusions

Using physiological and biochemical markers we were able to differentiate distinct phases during the annual cycle of grapevine bud growth, dormancy and bud bursts, something not possible using a phenological approach. At the cellular level, we demonstrated that the decrease of dormancy produced in late winter correlates with an increase in the proportion of cells in the G2 phase of the cell cycle and a decrease of cells in the G1 phase. At the molecular level, we observed that (i) the para-dormancy period is accompanied by greater levels of ADP-glucose, shikimate, tartrate, myo-inositol, pyruvate, and many TCA cycle intermediates; (ii) the period of endo-dormancy is characterized by the lowest levels of hexoses, their phosphorylated forms, and glycolytic intermediates; (iii) while the eco-dormancy period is characterized by greater levels of disaccharides such as sucrose (and Suc6P), and trehalose, along with monosaccharides, such as glucose and galactose, precursors and members of the raffinose-family oligosaccharides (e.g. galactinol and stachyose, plus the dicarboxylic acid malate. Thus, we conclude that the contents of different sugars in the grapevine buds have potential to be used as markers to differentiate dormancy and quiescent states. Finally, our data provides suggestive evidence that the role of Tre6P in promoting bud outgrowth can be a conserved mechanism in woody perennial plants.

## Acknowledgements

We thank Dr Karlia Meitha and Dr Wisam Salo for their help in this research. SS is an active member of the National System of Researchers (SNI, Sistema Nacional de Investigadores) and PEDECIBA from Uruguay. S.S. is supported by the CSIC I+D groups program (Uruguay), group name: “Food and Plant Biology” and group number “883431”. The authors acknowledge the facilities, and scientific and technical assistance of Microscopy Australia at the Centre for Microscopy, Characterisation & Analysis, The University of Western Australia. MC acknowledges funding support from the Australian Research Council (DP150103211; FT180100409). R.F. and J.E.L. were financially supported by the Max Planck Society.

